# Target-enriched metagenomics-informed qPCR detects rare, potentially dangerous β-lactamase genes in wastewater

**DOI:** 10.1101/2025.07.21.665820

**Authors:** Yuqing Mao, Mai Nguyen Tuan Anh, Kien Dang, Joanna L. Shisler, Thanh H. Nguyen

**Affiliations:** Department of Civil and Environmental Engineering, The Grainger College of Engineering, University of Illinois at Urbana-Champaign; Carl R. Woese Institute for Genomic Biology, University of Illinois at Urbana-Champaign; VinUni-Illinois Smart Health Center, University of Illinois at Urbana-Champaign; Department of Microbiology, University of Illinois at Urbana-Champaign

**Keywords:** Metagenomic sequencing, quantitative polymerase chain reaction, antimicrobial resistance, β-lactamase, primer design, wastewater

## Abstract

The rapid emergence of novel antibiotic resistance genes (ARGs) decreases the effectiveness of empirical antibiotic treatment for pathogen infections. Environmental ARG surveillance is an early-warning approach that can better inform antibiotic usage. Quantitative polymerase chain reaction (qPCR) is widely used for ARG surveillance because qPCR is easy, fast, and highly sensitive. However, it can only identify DNA targeted by pre-selected primers. In contrast, metagenomic sequencing can identify ARGs agnostically. However, sequencing is more expensive, less sensitive, and requires lengthy analysis of complex data sets. Target-enrichment metagenomic sequencing (TEMS), a method we developed previously, can mitigate the disadvantage of low sensitivity of traditional metagenomic sequencing. In this study, we propose a hybrid ARG surveillance pipeline for wastewater that capitalizes on both TEMS and qPCR. It uses a large-scale target determination of ARGs by using TEMS, followed by fine-scale routine surveillance of ARGs using qPCR. To connect the two steps, we developed a primer design tool, MSEDAP, which automatically analyzes metagenomic sequencing data and designs qPCR primers for ARG surveillance. This new workflow was ground-truthed using wastewater samples. TEMS identified seventeen β-lactamase gene targets of potential clinical importance. qPCR validated their presence and abundance using primers generated by MSEDAP.

**Synopsis:** A metagenomics-qPCR hybrid scheme can be used for environmental surveillance of antibiotic resistance genes (ARGs) to discover emerging ARGs originating from communities.

**Graphical abstract:** 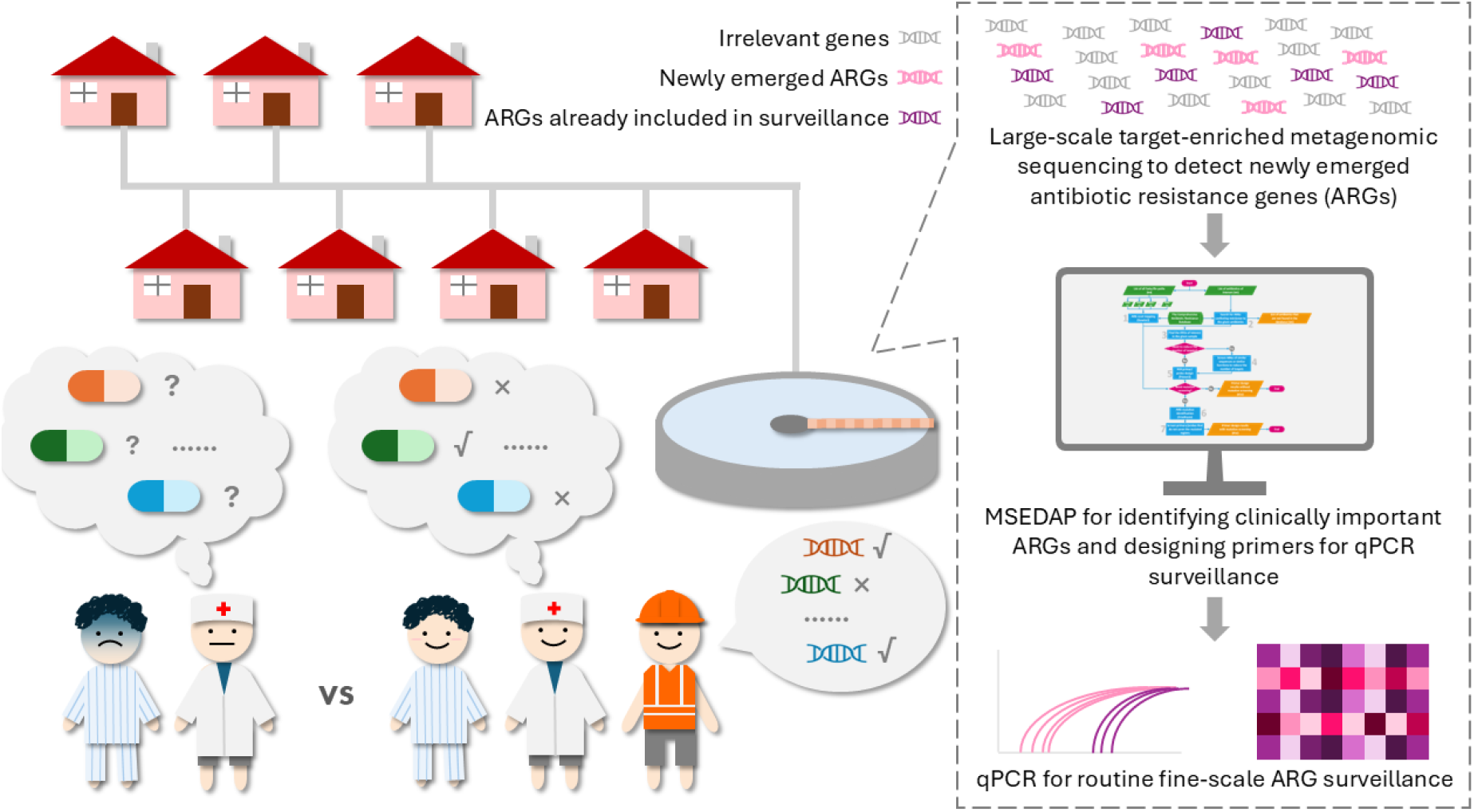

## 1 Introduction

The global spread of antimicrobial resistance (AMR) reduces the effectiveness of antibiotics in treatment of pathogen infections.^1^ Novel antibiotic resistance genes (ARGs) are emerging rapidly in addition to the spread of well-known ARGs. For example, the number of AMR gene families in the Comprehensive Antibiotic Resistance Database (CARD) increased from 304 to 458 from 2020-2023. ^2,3^ In hospitals and other clinical settings, the choice of antibiotic use is often based on common ARGs, including some β-lactamase genes (e.g., KPC, VIM, and OXA-48-like). ^4^ This approach is effective at responding quickly to current AMR threats but does not allow public health officials to identify emerging AMR threats outside of clinical settings.

The environmental surveillance of AMR can act as an early-warning system for introduction of new ARGs into a community. ^5^ It can provide population-level AMR prevalence, including ARGs that may not yet be present in hospitals and clinics. ^6^ A well-known example of ARG dissemination from the environment to clinics is the colistin resistance gene *mcr-1* that was originally found in commensal *Escherichia coli* in food animals. ^7^ Commonly used environmental ARG detection methods are quantitative polymerase chain reaction (qPCR) and metagenomic sequencing. ^8,9^ The advantages of qPCR include simple protocols, fast turnaround time, and high sensitivity, but qPCR requires the pre-determination of ARG targets for primer design and validation. In contrast, metagenomic sequencing does not require primer design, making it possible to detect a wide range of known and unknown ARGs. However, the high cost, long turnaround time, complex protocols, and the relatively low sensitivity of sequencing hinder the wide application of metagenomics in routine ARG surveillance.

There is a need for an easy-to-use, fast ARG surveillance system that capitalizes on metagenomic sequencing results for qPCR-based quantification of ARGs.^8,9^ Under such a pipeline, results from metagenomic sequencing would rapidly and precisely inform primer design for qPCR surveillance of environmental ARGs. Recently, we developed a CRISPR-based target-enrichment metagenomic sequencing (TEMS) method for ARG enrichment and detection.^10^ This method provides a 10-fold higher sensitivity to identify ARG targets, mitigating the normal drawback of low sensitivity of metagenomic sequencing. Unfortunately, the process going from metagenomic sequencing data analysis to qPCR primer design for ARGs remains cumbersome. To reduce the complexity of connecting metagenomics and qPCR primer design, we developed a primer design tool, MSEDAP (<u>M</u>etagenomic <u>Se</u>quencing <u>D</u>ata to <u>A</u>RG <u>P</u>rimers), that determines ARG targets for qPCR-based routine surveillance. MSEDAP generates primers and probes for qPCR-based ARG surveillance by using the metagenomic sequencing raw reads and a list of antibiotics in current use as inputs. With MSEDAP, the complexity of the in-silico pipeline between metagenomic sequencing and qPCR can be reduced to one command, making its use straightforward and easy for public health facilities. We tested this new pipeline by probing β-lactamase genes in urban wastewater collected over a six-month period. β-lactam antibiotics account for 65% of the current antibiotic market.^11^ Though many of the β-lactamase genes recorded in databases haven’t attracted attention in clinical settings, their mobility and resistance to clinically-important antibiotics are causing public health concern.^12^ These genes, especially metallo-β-lactamase genes, may greatly reduce the treatment choices for bacterial infection. ^13,14^ The aim of this study is to propose a wastewater ARG surveillance scheme that can provide real-time updates to healthcare facilities about the newly emerged AMR risks that arise from communities.

## 2 Methods

### 2.1 Sample collection, pre-processing, and DNA extraction

A total of 22 wastewater samples were collected from five wastewater treatment plants (WWTPs) in Chicago from June 7th to November 28th in 2023. The wastewater samples were transported to the lab on the day of collection in coolers filled with ice packs. The samples were stored at 4^°^C for a maximum of three days before pre-processing. For pre-processing, 20-75 mL of each well-mixed wastewater sample was filtered through 0.45 μm mixed cellulose esters (MCE) membrane filters until the filters clogged. For every sampling batch, a blank sample was created by filtering 50 mL of nuclease-free water in parallel. After filtering, all filters were stored at -80^°^C until DNA extraction.

For each sample, half of the filter was used for DNA extraction with FastDNA™ SPIN Kit for Soil (MP Biomedicals). The extraction steps followed the user manual with a minor modification: the duration of the centrifugation step after the bead-beating step was increased to 15 minutes to separate liquid and solid phases more thoroughly. Each DNA sample was eluted with 100 μL of the DES Elution Solution. DNA concentrations were quantified by Qubit (Thermo Fisher). The DNA samples were stored at -20^°^C until further analysis. Detailed information on sampling sites, volumes used for filtration, and DNA concentrations are in **Table S1**.

### 2.2 Metagenomics-informed qPCR primer design by MSEDAP

The command-line primer design tool MSEDAP can be found on GitHub page: https://github.com/Nguyen205/MSEDAP-Metagenomic-Sequencing-Data-to-ARG-Primers.git. Two inputs were required by the tool: a list of metagenomic sequencing fastq files and a list of antibiotic names we obtained from state of Illinois public health authorities. The sequencing list contained six paired-read fastq files which were retrieved from NCBI BioProject PRJNA1148079. ^10^ Cephalosporin, carbapenem, and monobactam were used as the antibiotic input, and these were chosen based on input from the Public Health Department for the State of Illinois. Any ARGs with gene mapping coverage values greater than 70% were considered present in the sample. Among these ARGs, MSEDAP screened out those that confer resistance to the antibiotics in the input list. All ARGs in CARD had been pre-grouped by their similarities determined by calculating pairwise average nucleotide identities (ANIs) before executing MSEDAP. The ARGs with ANIs and coverages greater than 90% were divided into the same group. One random ARG was selected from each group for primer design. Ten primers were designed for each ARG target. The qPCR amplicon lengths were set to the range of 75-150. To ensure the best primer binding efficiency for local surveillance, an additional mutation screening step was conducted to exclude the primer sets that covered the mutated sites in the corresponding ARGs detected in the wastewater samples. The complete primer design results can be found in the “publication_data” directory on the GitHub page.

### 2.3 qPCR for ARG detection

After primer design by MSEDAP, we selected 17 primer sets to validate their presence and abundance in different wastewater samples by qPCR. The primers targeting GES-26 were obtained from a previous testing run of MSEDAP, while the remaining primers were obtained in the final run. The primer specificity was cross-checked by NCBI Primer-BLAST. ^15^ Each qPCR assay had a total volume of 15 μL, including 1 μL of DNA template, 400 nM of each forward and reverse primer (**Table S2**), and 7.5 μL PowerUp™ SYBR™ Green Master Mix for qPCR (Applied Biosystems). The DNA samples were 10-fold diluted before loading to the plate to avoid PCR inhibition. All samples were loaded in triplicates. The thermal cycle consisted of a hold stage of 2 min at 50 °C and 10 min at 95 °C, a PCR stage with 40 cycles of 10 s at 95 °C and 1 min at 60 °C. A melt curve stage was performed after the PCR stage to identify the non-specific amplification. For each primer set in each qPCR run, triplicate non-template controls were created by replacing the 1 μL DNA sample with 1 μL nuclease-free water. All qPCRs were performed with 0.2 mL 96-well plates in QuantStudio™ 3 and QuantStudio™ 7 Real-Time qPCR Systems (Thermo Fisher). The qPCR performed in this study followed MIQE guidelines (**Table S3**).^16^

### 2.4 Primer efficiency validation

Primer efficiencies and sample inhibition levels were determined by making calibration curves.^17^ The DNA template for making calibration curves was made by pooling equal volumes of wastewater samples collected in this study, except for VIM-43 whose calibration curve was made by a synthesized template. Then, the DNA template was serial diluted by two or three-fold to create calibration curves. The different dilution levels of the DNA template were loaded in triplicates for qPCR. The same qPCR protocol was used in this step as described above. The detailed primer efficiency validation results are provided in **Table S2**.

### 2.5 Statistical analysis

The qPCR results were retrieved from Design & Analysis Software 2.8.0 (Thermo Fisher). The Cq value of an ARG in a sample was determined by averaging the Cq values of the triplicates. The outlier was defined as the replicate that differed from the other two replicates by at least 1 Cq. When more than two replicates showed negative signals, the corresponding ARG was determined as not detected in the sample. Contamination was negligible if the average Cq value of the non-template control of an ARG was greater than the Cq values of any DNA samples in the same run by 5 or more.

The relative abundances of the ARGs were determined by the ΔCq method. The 16S rRNA gene was used as the reference gene. The equations for calculating the relative abundances are listed below:

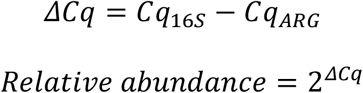

In the equations above, Cq_16S_ means the average Cq value of the 16S rRNA gene detected in each sample, and Cq_ARG_ means the average Cq value of the corresponding ARG detected in each sample.

To determine the lower limit of relative abundance quantification, we made a standard curve using a synthesized DNA template of 16S rRNA gene with known copy numbers. A theoretical Cq value of 34.995 corresponding to one copy of 16S rRNA gene per reaction was determined by the standard curve. Any ARGs with Cq values greater than this threshold were determined absent in the corresponding sample.

With the relative abundances, the app “Heatmap with Dendrogram” in Origin 2024b was used to plot heatmaps and perform hierarchical clustering among the ARGs and among wastewater samples. Euclidean distances were calculated among ARGs and among sampling sites. The dendrograms were clustered by the Ward’s method.

## 3 Results and Discussion

### 3.1 Code output for the MSEDAP algorithm to design primers

The MSEDAP algorithm includes seven steps. First, all fastq files containing the metagenomic sequencing results in the input list will be aligned to the reference ARG sequences obtained from CARD to generate a list of ARGs detected in the given samples. Second, with the list of antibiotics of current use, MSEDAP searches CARD and summarizes all ARGs conferring resistance to these antibiotics. Third, according to the ARG alignment results, the ARGs that are not present in the given sample will be excluded from primer design. Fourth, according to the qPCR capabilities, the user can select whether to exclude part of the ARG targets that share highly similar sequences or similar functions. Fifth, after all excluding steps, the ARG targets will be transferred to Primer3 for primer and probe design.^18^ If the mismatches in primer binding need to be considered, there will be two more optional steps, otherwise a summary of all primers and probes designed in step 5 will be the final output. Step 6 uses FreeBayes to summarize all mutations in the ARGs detected in the given sample, and step 7 excludes all primers and probes that cover the mutated sites from the output of step 5 to ensure the best amplification efficiency.

### 3.2 Rare β-lactamase genes were detected in WWTPs across Chicago area

The robustness of the proposed method was evaluated by probing wastewater samples for ARGs that confer resistance to cephalosporin, carbapenem, and monobactam. Using the CRISPR-enriched metagenomics method to identify ARGs, 17 β-lactamase genes were successfully detected by qPCR in 22 samples collected from five WWTPs in the greater Chicago area over a six-month period. ^10^ Most global surveillance strategies monitor for five groups of β-lactamase genes: VIM, NDM, IMP, KPC, and OXA-48-like. ^19^ Interestingly, TEMS only identified genes within three of these groups (VIM-43, KPC-59, and OXA-245) as present in Chicago-area wastewater. More importantly, this method identified the presence of 14 β-lactamase genes that are usually not monitored. We then used MSEDAP to design primers for all 17 β-lactamase genes. The newly-designed 17 primer sets showed adequate amplification efficiencies and high specificity in the primer validation step, with efficiencies ranging from 92.9% to 114.6%, R^2^ values greater than 0.95, and the non-template controls had Cq values of at least 5 greater than the Cq values of the lower limits of the linear dynamic range (**Table S2**).

When using qPCR, the relative abundances of different β-lactamase genes in samples varied from 0 to 1.48 × 10^−2^ (**Figure 2**). The relative abundance of 1.48 × 10^−2^ means there are 1.48 copies of an ARG in every 100 bacteria cells. The OXA-2 gene had the highest relative abundance (2.05 × 10^−3^ to 1.48 × 10^−2^) among the 17 ARGs targeted in this study. This suggests that there is a high prevalence of the OXA-2 gene in wastewater bacteria that can confer resistance to ceftazidime, amoxicillin, and ampicillin. Eight of the 17 β-lactamase genes (VIM-43, GPC-1, ACI-1, CMY-159, CTX-M-27, SHV-12, BEL-2, and AIM-1) were present at relative abundances that are considered low (below 10^−4^) in all samples. These low-abundance genes were found by TEMS but not by conventional metagenomic sequencing, ^10^ demonstrating that TEMS is more sensitive in identifying ARG targets than conventional metagenomic sequencing.

**Figure 1.**
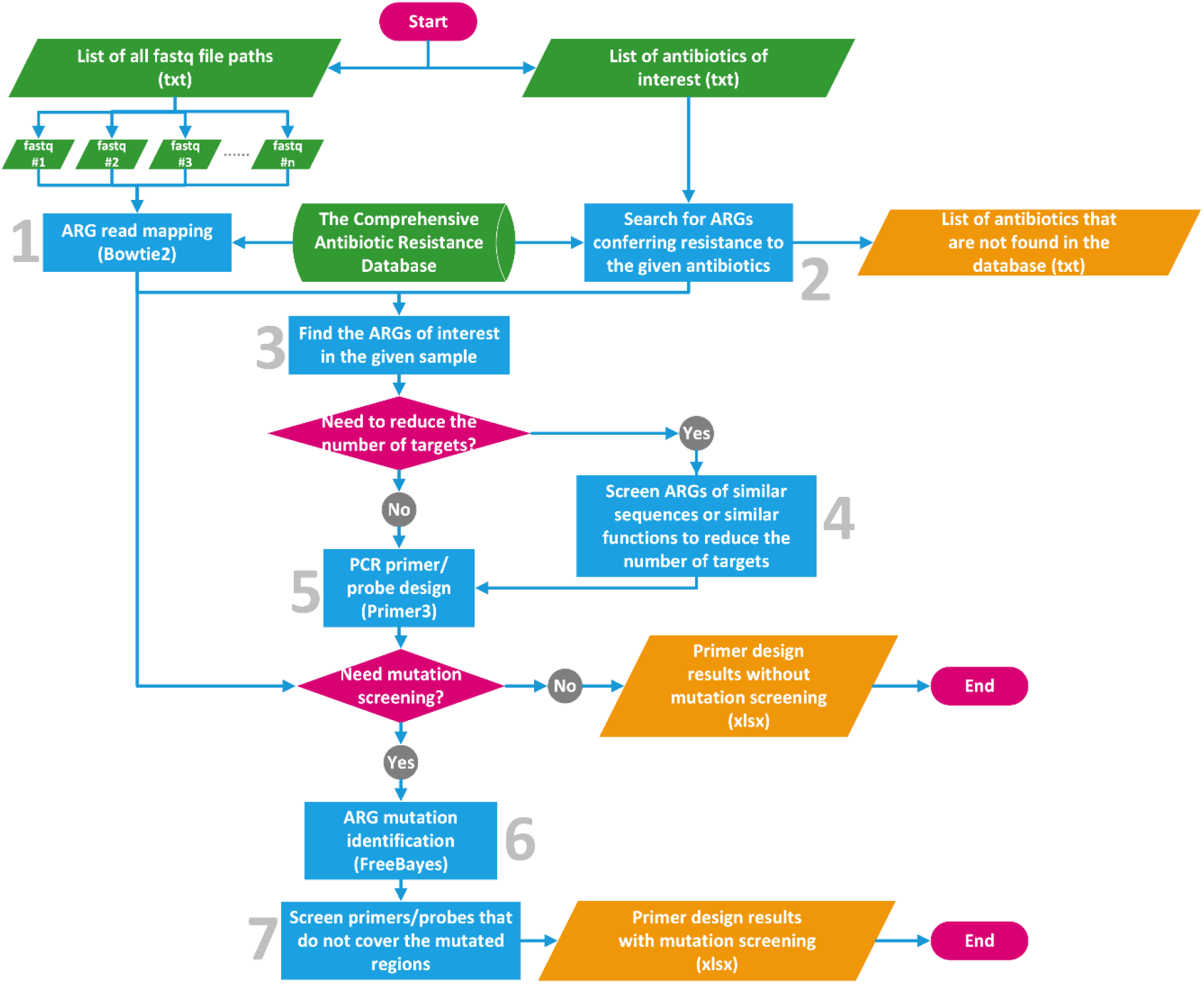
The algorithm flow chart of MSEDAP. The green parallelograms indicate the input files. The orange parallelograms indicate the output files. The green cylinder indicates the database. The blue rectangles indicate the processing steps. The pink diamonds indicate decision steps.

**Figure 2.**
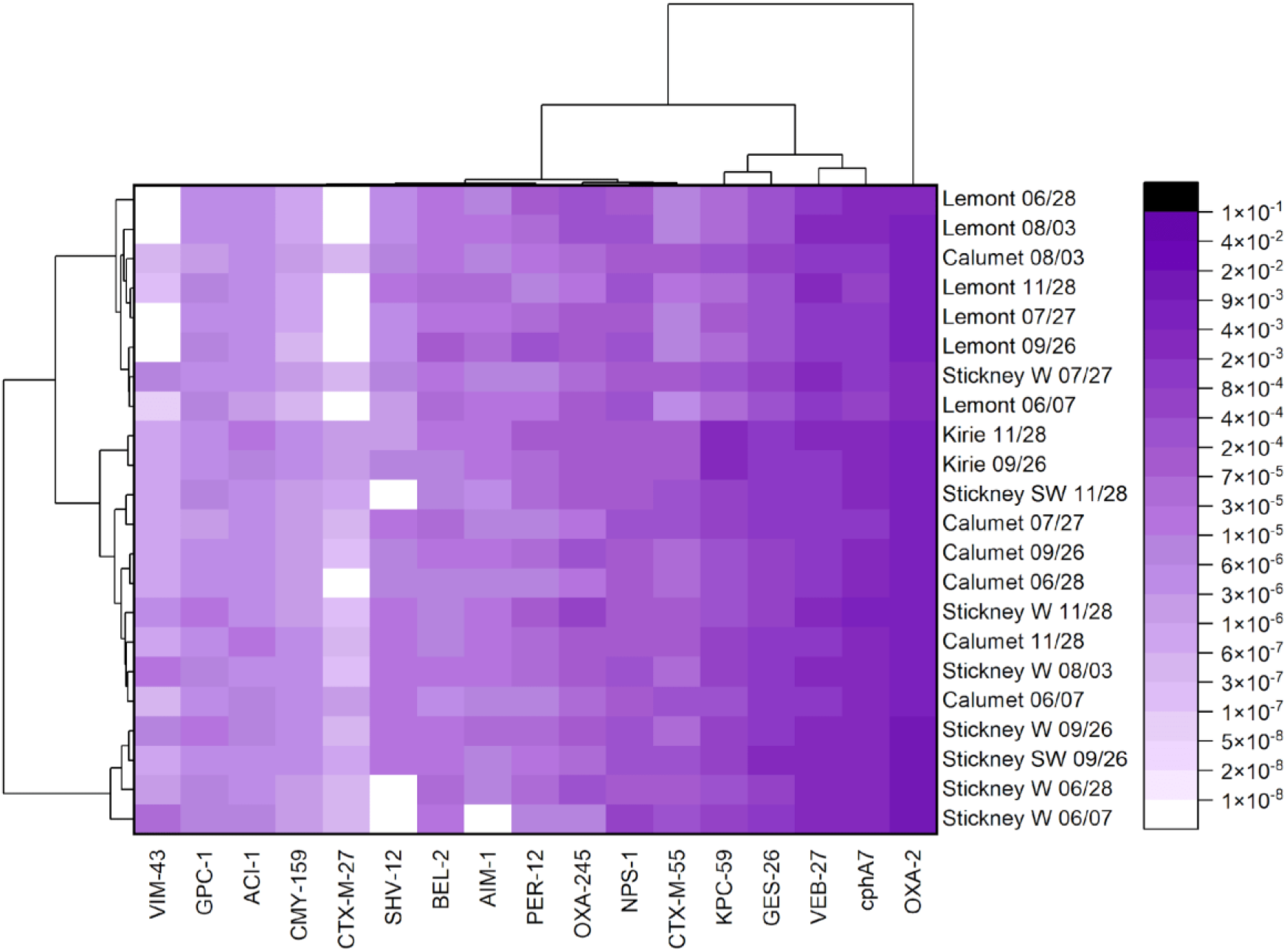
The heatmap of the 17 β-lactamase genes in 22 Chicago wastewater samples collected from five Chicago WWTPs: Calumet, Kirie, Lemont, Stickney W, and Stickney SW. The numbers next to the plants’ names are the month and days of sample collection during 2023. The legend shows the scale of the relative abundances of the targeted β-lactamase genes. The dendrogram on the left shows the clustering of different WWTPs, based on similarity in ARG distribution. The dendrogram on the top shows the clustering of the β-lactamase genes.

Most β-lactamase genes were detected consistently through the six-month sampling period in each WWTP (**Figure 2**). The ARGs present in samples from Lemont and Kirie, WWTPs that serve two small populations, formed two different clusters (**Figure 2; Figure S1**). The relative abundance ranges of KPC-59 (3.53 × 10^−5^ – 7.78 × 10^−5^ vs 2.82 × 10^−3^ – 3.59 × 10^−3^), CTX-M-55 (6.07 × 10^−6^ – 1.49 × 10^−5^ vs 7.62 × 10^−5^ – 1.40 × 10^−4^), and VIM-43 (0 – 1.92 × 10^−7^ vs 6.34 × 10^−7^ – 7.80 × 10^−7^) did not overlap between these two WWTPs. In contrast, WWTPs serving population sizes larger than one million (i.e., Calumet, Stickney W, and Stickney SW) showed more general ARG distribution patterns (**Figure 2; Figure S1**). The different ARG distributions in different WWTPs supported the necessity of conducting community-scale qPCR surveillance.

Different classes of β-lactamase genes possess different β-lactam degradation mechanisms, leading to different clinical treatment strategies. Three of the 17 β-lactamase genes targeted in this study (VIM-43, AIM-1, and *cphA7*) encode metallo-β-lactamases. Clinical treatment of infections caused by metallo-β-lactamase-producing pathogens is exceptionally challenging because metallo-β-lactamases are activated by metal ions and therefore cannot be inhibited by β-lactamase inhibitors.^20^ Current ARG surveillances often focus on B1 metallo-β-lactamases, such as IMP, NDM, and VIM, ^21^ because the corresponding ARGs have been identified in clinical settings. Another major subgroup of metallo-β-lactamases, B3 metallo-β-lactamases, were often considered immobile and hosted by commensal bacteria, making the corresponding ARGs lack attention. ^22^ However, in this study, in addition to VIM-43 (which already is included in the current ARG surveillance scheme), a B3 metallo-β-lactamase gene AIM-1 with high mobility that was originally hosted by *Pseudomonas aeruginosa* was detected in most wastewater samples (relative abundances up to 4.47 × 10^−5^). ^23–25^ This finding suggested the need for increased attention to B3 metallo-β-lactamases to better prepare for their future dissemination into a wider range of pathogenic bacteria.

## 4 Environmental relevance

Based on the findings in this study, we propose a scheme for wastewater-based ARG surveillance with improved efficiency that benefits from both metagenomic sequencing and qPCR. First, CRISPR-based target-enrichment metagenomic sequencing, TEMS, is conducted for wastewater samples on a large scale, for example, city-wide wastewater sequenced annually or semiannually. Samples from different WWTPs in the same city may be pooled and sequenced together. Second, after obtaining the large-scale ARG presence patterns by metagenomic sequencing, primers for local qPCR surveillance can be designed accordingly using tools like MSEDAP described in this study. Finally, with the designed primers, qPCR surveillance can be conducted on a finer scale, like community-scale weekly reports, for ARG risk early warning. This scheme can utilize resources more efficiently in ARG surveillance by identifying newly emerging ARGs and reducing the resources spent on tracking non-existent targets.

## Supporting information

<u>Figure S1</u>. The locations of the wastewater treatment plants (WWTPs) for sample collection in this study. Stickney W and Stickney SW are in the same location. The population sizes served by each WWTP are labeled in the parentheses below the WWTP names. Among the five WWTPs, the population sizes served by Calumet (1.13 M), Stickney A (1.13 M) and Stickney B (1.13 M) were greater than Lemont (13,098) and Kirie (270,647). (TIF)

<u>Table S1</u>. Detailed information about wastewater samples. (XLSX)

<u>Table S2</u>. Detailed information about primer amplification efficiency validation results. (XLSX)

<u>Table S3</u>. MIQE guideline checklist. (XLSX)

## Acknowledgement

We acknowledge funding from the Water Research Foundation (5182), NTU/U of I Joint Research and Innovation Seed Grants Program, and USEPA (R840487). This study has not been formally reviewed by EPA. EPA does not endorse any products or commercial services mentioned in this publication. We acknowledge Dr. Rebecca Lee Smith for working with the public health board. MNTA and KD acknowledged scholarships from VinUni Illinois Smart Health Center. We acknowledge the effort in sample collection from Arthur R. Schmidt IV, Ryan McAllister, Amaja Craft, and Josie Hoppenworth. We acknowledge Dr. Albert Cox, Dr. Essam El-Naggar, Dr. Kamlesh Patel, and Dr. Kaylyn Patterson from the Metropolitan Water Reclamation District of Greater Chicago for providing WWTP sewage samples. We acknowledge Roy J. Carver Biotechnology Center for its sequencing services.

## Abbreviations

AMR: antimicrobial resistance
ARG: antibiotic resistance gene
CARD: the Comprehensive Antibiotic Resistance Database
TEMS: target-enrichment metagenomic sequencing
qPCR: quantitative polymerase chain reaction
WWTP: wastewater treatment plant
MIQE: minimum information for publication of quantitative real-time PCR experiments

## References

(1) Zhang, S.; Yang, J.; Abbas, M.; Yang, Q.; Li, Q.; Liu, M.; Zhu, D.; Wang, M.; Tian, B.; Cheng, A. Threats across Boundaries: The Spread of ESBL-Positive Enterobacteriaceae Bacteria and Its Challenge to the “One Health” Concept. Front Microbiol 2025, 16, 1496716. 10.3389/FMICB.2025.1496716/XML/NLM.

(2) Alcock, B. P.; Raphenya, A. R.; Lau, T. T. Y.; Tsang, K. K.; Bouchard, M.; Edalatmand, A.; Huynh, W.; Nguyen, A. L. V.; Cheng, A. A.; Liu, S.; Min, S. Y.; Miroshnichenko, A.; Tran, H. K.; Werfalli, R. E.; Nasir, J. A.; Oloni, M.; Speicher, D. J.; Florescu, A.; Singh, B.; Faltyn, M.; Hernandez-Koutoucheva, A.; Sharma, A. N.; Bordeleau, E.; Pawlowski, A. C.; Zubyk, H. L.; Dooley, D.; Griffiths, E.; Maguire, F.; Winsor, G. L.; Beiko, R. G.; Brinkman, F. S. L.; Hsiao, W. W. L.; Domselaar, G. V.; McArthur, A. G. CARD 2020: Antibiotic Resistome Surveillance with the Comprehensive Antibiotic Resistance Database. Nucleic Acids Res 2020, 48 (D1), D517–D525. 10.1093/NAR/GKZ935.

(3) Alcock, B. P.; Huynh, W.; Chalil, R.; Smith, K. W.; Raphenya, A. R.; Wlodarski, M. A.; Edalatmand, A.; Petkau, A.; Syed, S. A.; Tsang, K. K.; Baker, S. J. C.; Dave, M.; Mccarthy, M. C.; Mukiri, K. M.; Nasir, J. A.; Golbon, B.; Imtiaz, H.; Jiang, X.; Kaur, K.; Kwong, M.; Liang, Z. C.; Niu, K. C.; Shan, P.; Yang, J. Y. J.; Gray, K. L.; Hoad, G. R.; Jia, B.; Bhando, T.; Carfrae, L. A.; Farha, M. A.; French, S.; Gordzevich, R.; Rachwalski, K.; Tu, M. M.; Bordeleau, E.; Dooley, D.; Griffiths, E.; Zubyk, H. L.; Brown, E. D.; Maguire, F.; Beiko, R. G.; Hsiao, W. W. L.; Brinkman, F. S. L.; Van Domselaar, G.; Mcarthur, A. G. CARD 2023: Expanded Curation, Support for Machine Learning, and Resistome Prediction at the Comprehensive Antibiotic Resistance Database. Nucleic Acids Res 2023, 51 (D1), D690–D699. 10.1093/NAR/GKAC920.

(4) Tamma, P. D.; Heil, E. L.; Justo, J. A.; Mathers, A. J.; Satlin, M. J.; Bonomo, R. A. Infectious Diseases Society of America 2024 Guidance on the Treatment of Antimicrobial-Resistant Gram-Negative Infections. Clinical Infectious Diseases 2024, 00. 10.1093/CID/CIAE403.

(5) Liguori, K.; Keenum, I.; Davis, B. C.; Calarco, J.; Milligan, E.; Harwood, V. J.; Pruden, A. Antimicrobial Resistance Monitoring of Water Environments: A Framework for Standardized Methods and Quality Control. Environ Sci Technol 2022, 56 (13), 9149–9160. 10.1021/ACS.EST.1C08918/ASSET/IMAGES/LARGE/ES1C08918_0005.JPEG.

(6) Conforti, S.; Pruden, A.; Acosta, N.; Anderson, C.; Buergmann, H.; Calabria De Araujo, J.; Cristobal, J. R.; Drigo, B.; Ellison, C.; Francis, Z.; Frigon, D.; Gaenzle, M.; Vierheilig, J.; Julian, T. R.; Klümper, U.; Ma, L.; Mangat, C.; Nadimpalli, M.; Nakashita, M.; Osena, G.; Rathinavelu, S.; Reid-Smith, R.; Saldana, M.; Schmitt, H.; Li, S.; Singer, A. C.; Tran, T. T.; Yanac, K.; Ybazeta, G.; Harnisz, M. Strengthening Policy Relevance of Wastewater-Based Surveillance for Antimicrobial Resistance. Environ Sci Technol 2025. 10.1021/ACS.EST.4C09663.

(7) Liu, Y. Y.; Wang, Y.; Walsh, T. R.; Yi, L. X.; Zhang, R.; Spencer, J.; Doi, Y.; Tian, G.; Dong, B.; Huang, X.; Yu, L. F.; Gu, D.; Ren, H.; Chen, X.; Lv, L.; He, D.; Zhou, H.; Liang, Z.; Liu, J. H.; Shen, J. Emergence of Plasmid-Mediated Colistin Resistance Mechanism MCR-1 in Animals and Human Beings in China: A Microbiological and Molecular Biological Study. Lancet Infect Dis 2016, 16 (2), 161–168. 10.1016/S1473-3099(15)00424-7.

(8) Taylor, W.; Bohm, K.; Dyet, K.; Weaver, L.; Pattis, I. Comparative Analysis of QPCR and Metagenomics for Detecting Antimicrobial Resistance in Wastewater: A Case Study. BMC Res Notes 2025, 18 (1), 1–6. 10.1186/S13104-02407027-9/FIGURES/2.

(9) Elbait, G. D.; Daou, M.; Abuoudah, M.; Elmekawy, A.; Hasan, S. W.; Everett, D. B.; Alsafar, H.; Henschel, A.; Yousef, A. F. Comparison of QPCR and Metagenomic Sequencing Methods for Quantifying Antibiotic Resistance Genes in Wastewater. PLoS One 2024, 19 (4), e0298325. 10.1371/JOURNAL.PONE.0298325.

(10) Mao, Y.; Shisler, J. L.; Nguyen, T. H. Enhanced Detection for Antibiotic Resistance Genes in Wastewater Samples Using a CRISPR-Enriched Metagenomic Method. Water Res 2025, 274, 123056. 10.1016/J.WATRES.2024.123056.

(11) Pandey, N.; Cascella, M. Beta-Lactam Antibiotics. StatPearls 2023.

(12) Castanheira, M.; Simner, P. J.; Bradford, P. A. Extended-Spectrum β-Lactamases: An Update on Their Characteristics, Epidemiology and Detection. JAC Antimicrob Resist 2021, 3 (3). 10.1093/JACAMR/DLAB092.

(13) Mojica, M. F.; Rossi, M. A.; Vila, A. J.; Bonomo, R. A. The Urgent Need for Metallo-β-Lactamase Inhibitors: An Unattended Global Threat. Lancet Infect Dis 2022, 22 (1), e28–e34. 10.1016/S1473-3099(20)30868-9/ASSET/7B28004F-9A96-4C06-8BC2-233C15BE0B3B/MAIN.ASSETS/GR2.JPG.

(14) Zakhour, J.; El Ayoubi, L. W.; Kanj, S. S. Metallo-Beta-Lactamases: Mechanisms, Treatment Challenges, and Future Prospects. Expert Rev Anti Infect Ther 2024, 22 (4), 189–201. 10.1080/14787210.2024.2311213;WGROUP:STRING:PUBLICATION.

(15) Ye, J.; Coulouris, G.; Zaretskaya, I.; Cutcutache, I.; Rozen, S.; Madden, T. L. Primer-BLAST: A Tool to Design Target-Specific Primers for Polymerase Chain Reaction. BMC Bioinformatics 2012, 13, 134. 10.1186/1471-2105-13-134.

(16) Bustin, S. A.; Benes, V.; Garson, J. A.; Hellemans, J.; Huggett, J.; Kubista, M.; Mueller, R.; Nolan, T.; Pfaffl, M. W.; Shipley, G. L.; Vandesompele, J.; Wittwer, C. T. The MIQE Guidelines: Minimum Information for Publication of Quantitative Real-Time PCR Experiments. Clin Chem 2009, 55 (4), 611–622. 10.1373/CLINCHEM.2008.112797.

(17) Mao, Y.; Akdeniz, N.; Nguyen, T. H. Quantification of Pathogens and Antibiotic Resistance Genes in Backyard and Commercial Composts. Science of The Total Environment 2021, 797, 149197. 10.1016/J.SCITOTENV.2021.149197.

(18) Untergasser, A.; Cutcutache, I.; Koressaar, T.; Ye, J.; Faircloth, B. C.; Remm, M.; Rozen, S. G. Primer3—New Capabilities and Interfaces. Nucleic Acids Res 2012, 40 (15), e115–e115. 10.1093/NAR/GKS596.

(19) Nordmann, P.; Poirel, L. Epidemiology and Diagnostics of Carbapenem Resistance in Gram-Negative Bacteria. Clinical Infectious Diseases 2019, 69 (Supplement_7), S521–S528. 10.1093/CID/CIZ824.

(20) Boyd, S. E.; Livermore, D. M.; Hooper, D. C.; Hope, W. W. Metallo-β-Lactamases: Structure, Function, Epidemiology, Treatment Options, and the Development Pipeline. Antimicrob Agents Chemother 2020, 64 (10), e00397–20. 10.1128/AAC.00397-20.

(21) Mentasti, M.; Prime, K.; Sands, K.; Khan, S.; Wootton, M. Rapid Detection of IMP, NDM, VIM, KPC and OXA-48-like Carbapenemases from Enterobacteriales and Gram-Negative Non-Fermenter Bacteria by Real-Time PCR and Melt-Curve Analysis. European Journal of Clinical Microbiology and Infectious Diseases 2019, 38 (11), 2029–2036. 10.1007/S10096-019-03637-5,.

(22) Krco, S.; Davis, S. J.; Joshi, P.; Wilson, L. A.; Monteiro Pedroso, M.; Douw, A.; Schofield, C. J.; Hugenholtz, P.; Schenk, G.; Morris, M. T. Structure, Function, and Evolution of Metallo-β-Lactamases from the B3 Subgroup—Emerging Targets to Combat Antibiotic Resistance. Front Chem 2023, 11, 1196073. 10.3389/FCHEM.2023.1196073/ENDNOTE.

(23) Amsalu, A.; Sapula, S. A.; Whittall, J. J.; Hart, B. J.; Bell, J. M.; Turnidge, J.; Venter, H. Worldwide Distribution and Environmental Origin of the Adelaide Imipenemase (AIM-1), a Potent Carbapenemase in Pseudomonas Aeruginosa. Microb Genom 2021, 7 (12), 000715. 10.1099/MGEN.0.000715/CITE/REFWORKS.

(24) Bougnom, B. P.; Zongo, C.; McNally, A.; Ricci, V.; Etoa, F. X.; Thiele-Bruhn, S.; Piddock, L. J. V. Wastewater Used for Urban Agriculture in West Africa as a Reservoir for Antibacterial Resistance Dissemination. Environ Res 2019, 168, 14–24. 10.1016/J.ENVRES.2018.09.022.

(25) Zhou, H.; Guo, W.; Zhang, J.; Li, Y.; Zheng, P.; Zhang, H. Draft Genome Sequence of a Metallo-β-Lactamase (BlaAIM-1)-Producing Klebsiella Pneumoniae ST1916 Isolated from a Patient with Chronic Diarrhoea. J Glob Antimicrob Resist 2019, 16, 165–167. 10.1016/J.JGAR.2019.01.010.

