## Supplementary figures and images for "Target-enriched metagenomics-informed qPCR detects rare, potentially dangerous β-lactamase genes in wastewater"

### Figure_S1_Map.tif

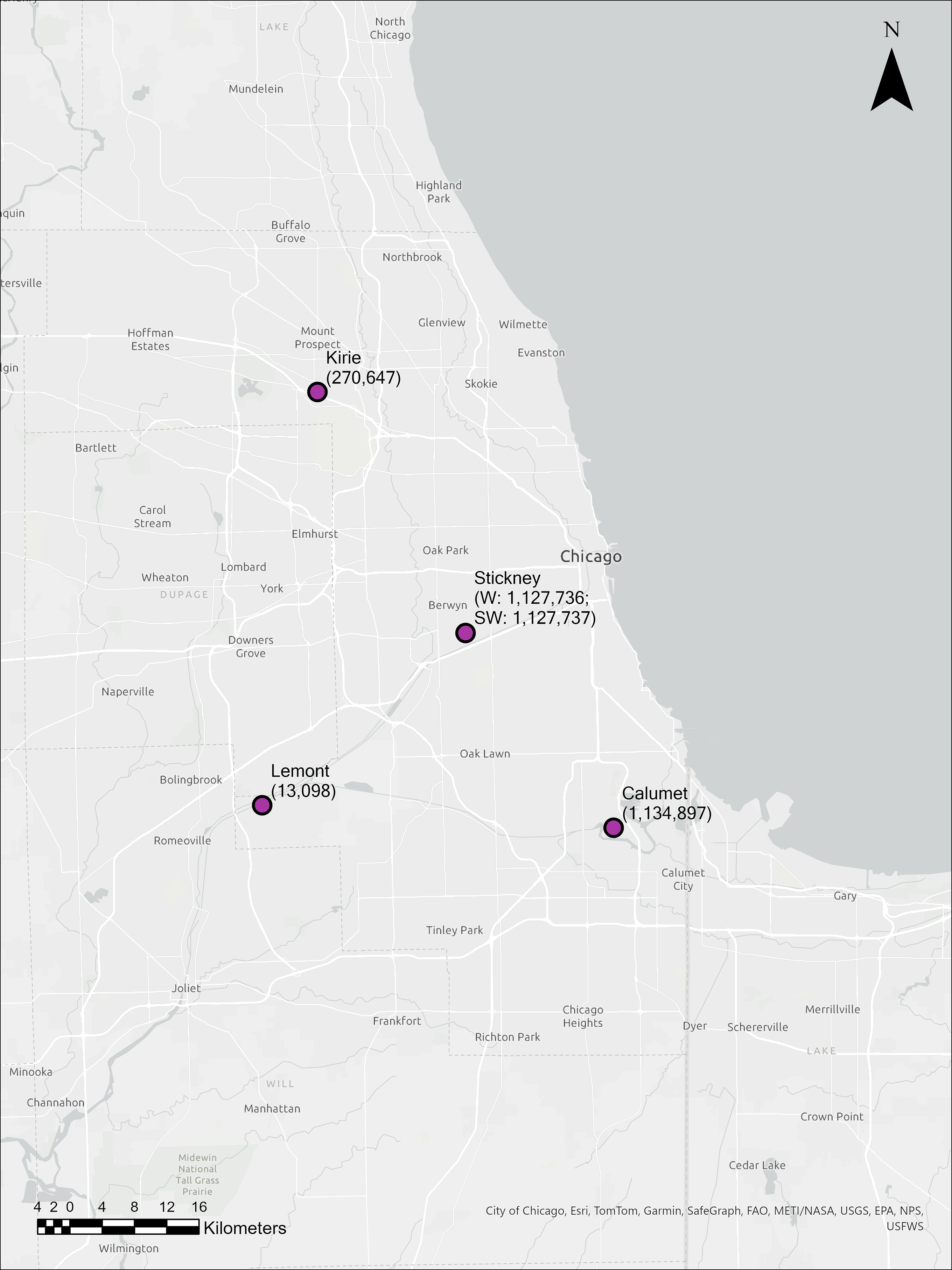
